# Hippocampal and neocortical oscillations are tuned to behavioral state in freely-behaving macaques

**DOI:** 10.1101/552877

**Authors:** Omid Talakoub, Patricia Sayegh, Thilo Womelsdorf, Wolf Zinke, Pascal Fries, Christopher M. Lewis, Kari L. Hoffman

## Abstract

Wireless recordings in macaque neocortex and hippocampus showed stronger theta oscillations during early-stage sleep than during alert volitional movement including walking. In contrast, hippocampal beta and gamma oscillations were prominent during walking and other active behaviors. These relations between hippocampal rhythms and behavioral states in the primate differ markedly from those observed in rodents. Primate neocortex showed similar changes in spectral content across behavioral state as the hippocampus.

## Main

In studies of human and non-human primates, the predominance of computer tasks and other stationary experiments has limited our understanding of neural changes across behavioral states, particularly for volitional, self-movement-related behaviors. By contrast, in rats, free behavior has been the predominant model approach, revealing well-established brain-behavior state dichotomies. For example, within rat hippocampus, theta-band oscillations appear consistently during locomotion, but also during other active voluntary movements and in REM sleep (‘Type II behaviors’). This activity appears in opposition to ripple-containing Large Irregular Activity (‘LIA’), which is seen during consummatory behaviors, self-grooming and in non-REM sleep (‘Type I behaviors’) ^1,2^. Rat hippocampal spectral tuning with behavioral state differs from the pattern observed in the neocortex, with the latter typically showing a prevalence of beta- and gamma-frequency activity during the greatest levels of vigilance and activity ^3–5^. These spectral shifts with behavior, however, may not be conserved across species ^3,6–8^. In macaques and humans engaged in stationary tasks, hippocampal theta can surprisingly be weaker during the ‘active’ states, with beta- and gamma-band power more closely coupled to active performance, similar to neocortical tuning ^9–16^. Results from ambulating humans suggest this may have been due to stationarity ^17,18^, raising the issue of how hippocampal and neocortical oscillations change across sleep and waking vigilance states in freely-moving primates.

To address this, we recorded a total of 210 hours of behaviors (supplementary Table 1) from three macaques who had chronically-implanted intracortical multichannel electrode arrays in the hippocampus (n=3 animals, Fig 1a), medial prefrontal and retrosplenial/posterior cingulate regions (n=2 animals). Behaviors were divided into 4 groups of increasing levels of activity/arousal (see Online Methods): (i) sleep; (ii) inactive (‘Type I’) states such as eating, drinking and being groomed; (iii) active (‘Type II’) including orienting responses, grooming another monkey, foraging, and (iv) walking (see supplementary video, Fig 1b). In both primate hippocampus and neocortex, we observed characteristic rhythms: A theta rhythm peaking around 7 Hz, a beta rhythm peaking around 12 Hz during sleep and around 12-18 Hz during waking depending on the region, and a hippocampal high-beta/low-gamma rhythm peaking broadly around 30 Hz and a cingulate gamma peaking more sharply at 46 Hz. Whereas activity in neocortex was generally consistent with that described in the rat ^3–5^, activity in hippocampus clearly showed different associations to behavioral states. For example, in the early stages of sleep, all sites showed prominent enhancements in theta-band (5–10 Hz) power compared to that of waking behaviors (Fig. 1c, sp<0.01 ranksum test on 1–10 Hz bands). Theta-band peaks were greater during sleep compared to waking, as measured relative to the 1/f component of the whole-session power spectrum (Fig 1c, left column). In contrast, hippocampal power in the beta and gamma bands was greater during activity than sleep (Fig. 1c, left column, p<0.01 ranksum test on 20–50 Hz bands). Power in all tested frequency bands between 10 and 80 Hz did not differ between inactive (‘Type I’) states and active Type II behaviors including walking (p>0.05, ranksum test).

**Fig 1.**
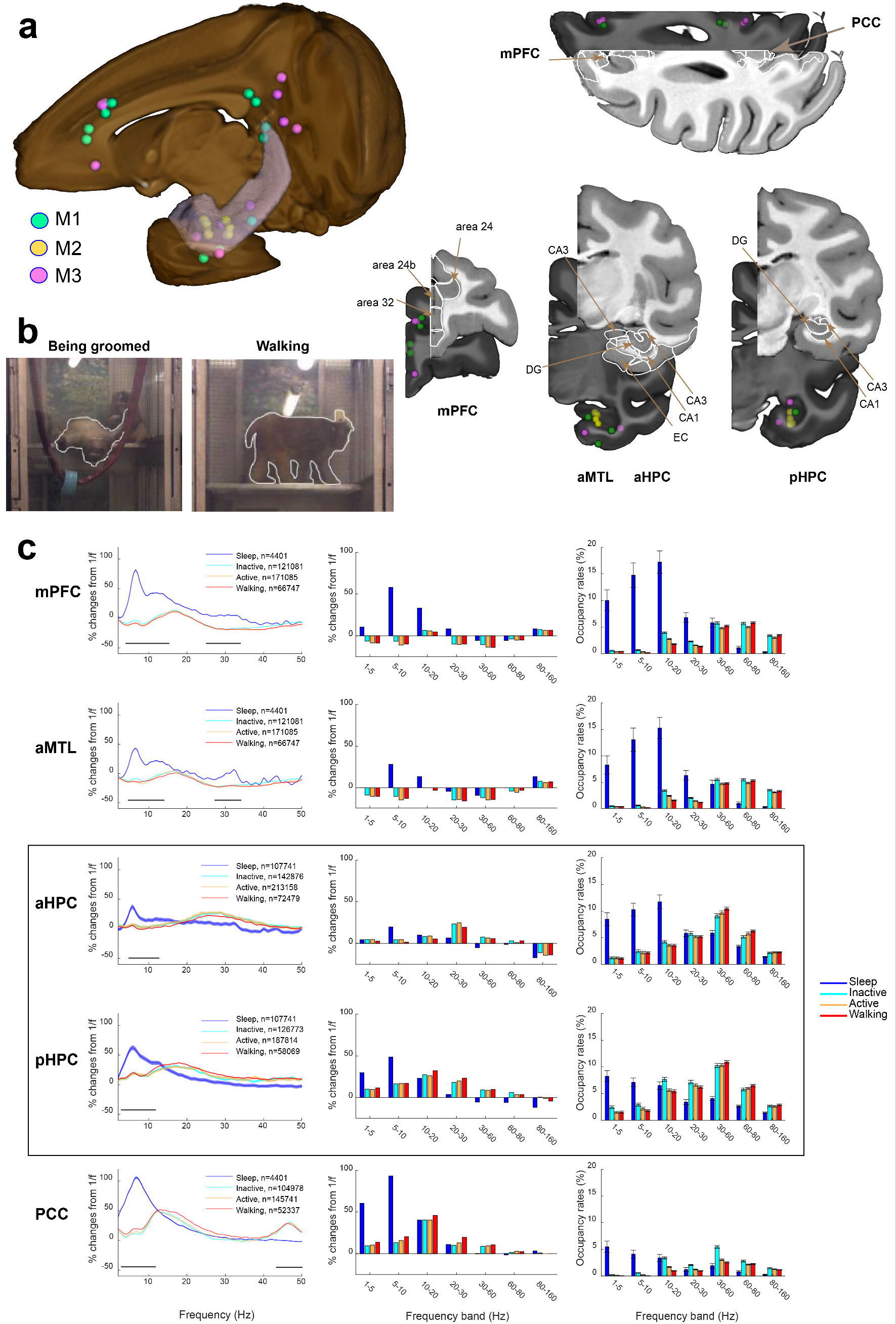
Spectral content recorded wirelessly across macaque hippocampal and neocortical sites, grouped by behavioral state. **a)**Electrode locations across animals were coregistered and marked on a cutaway of a macaque brain for Animal 1 in green, Animal 2 in yellow and Animal 3 in pink. Animal 2 electrode locations were projected onto the left hemisphere for comparison across animals. mPFC and MTL electrodes sites are shown projected onto coronal slices and PCC on an axial slice. b) Examples of video frames when Animal 1 was groomed in left and walking in right (see Supplementary Video). **c)**Frequency spectral analysis of mPFC, aMTL, aHPC, pHPC, and PCC regions during Sleep, Inactive, Active and Walking states (see online Methods). **Left)**Average FFT for each behavioral state measured relative to the 1/f normalized power over the full recording sessions, with 95% confidence boundaries shaded. Differences between sleep and waking power changes are indicated by the horizontal black lines. The number of 1-second epochs for each behavior is stated in the legends for each recording site, along with the color correspondence for each of the four behaviors. **Middle)**Percentile changes from 1/f normalized power for each frequency band. **Right)**Percentage of time (occupancy rate) of detected band-specific oscillations for each behavioral category. Error bars indicate standard deviations. Colormap shown on the far right.

By detecting band-limited bursts of spectral power, we could estimate the prevalence of oscillations across behavioral states. No single band of oscillations occurred more than 12% of the time the animal spent in any waking behavioral state. The greatest rates observed during waking were for the 30–60 Hz gamma band in the hippocampus. In contrast, hippocampal theta oscillations were rare during walking (~2%); they occurred roughly four times more often in sleep as in waking behavior (Fig. 1c, right column, t-test, p<0.01; normalized by the amount of time spent in each behavioral state). Hippocampal theta oscillations contained a greater proportion of long theta bouts during sleep compared to walking (p<0.001, KS-test, Fig. 2a,c). In contrast, long bouts of gamma (>30 Hz) were more common in hippocampus during active behavior compared with sleep (p<0.01, KS-test). Taken together, these results indicate the prevalence, magnitude, and duration of oscillations for a given set of behaviors, but not the inverse i.e. the likelihood of a behavioral state given the appearance of an oscillation.

**Fig 2.**
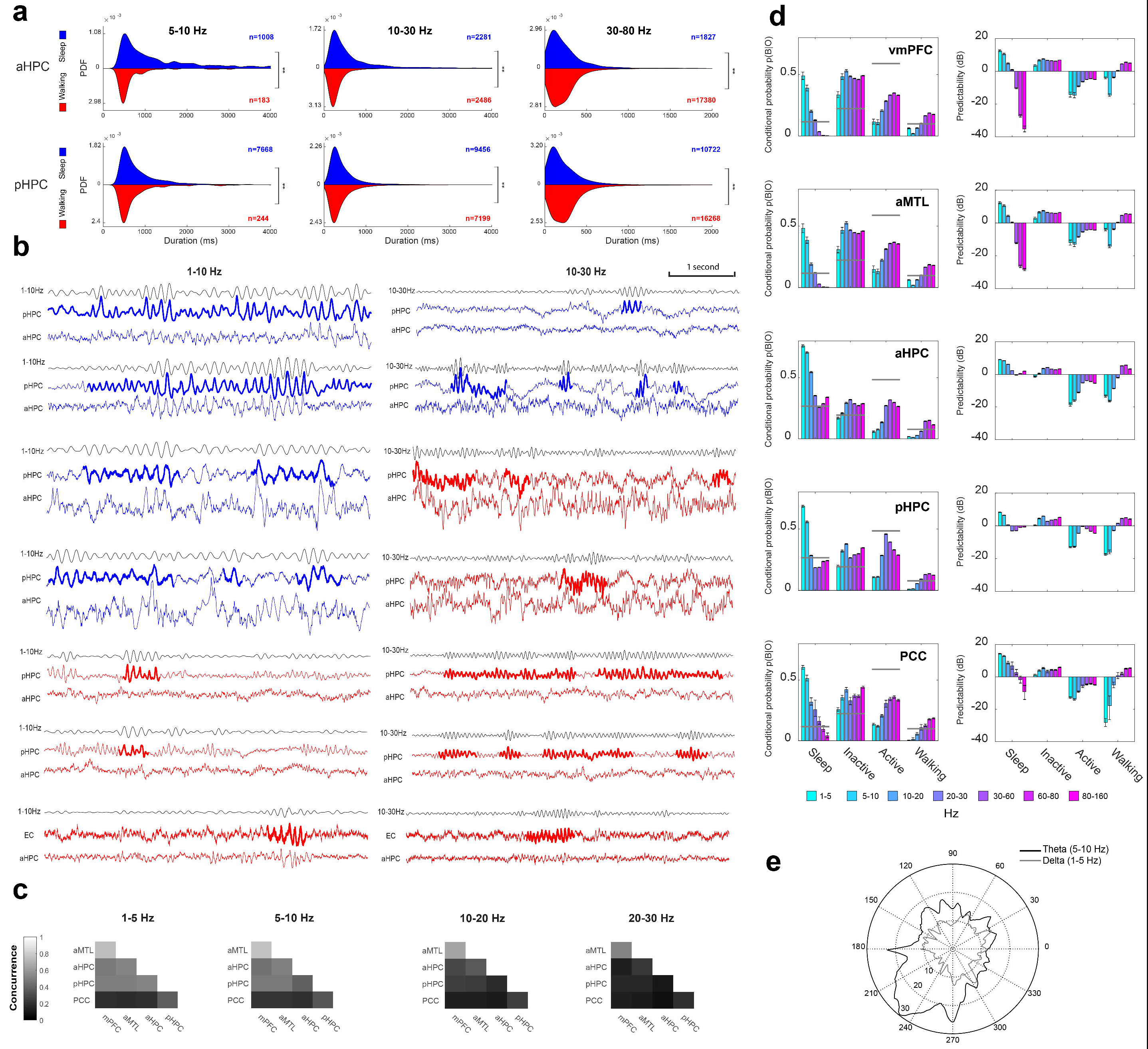
Oscillation durations and predictability of behavior given oscillations **a)**Distribution of oscillation durations for anterior and posterior hippocampal contacts during Sleep (blue) and Walking (red). X-axes are limited to 4 s for theta and beta bands, and to 2 s for gamma, to aid visibility of predominant durations. **b)**Examples for the detection of oscillations at 1–10 Hz (left) and 10–30 Hz (right) during 5 seconds of a continuous behavioral state. Broadband filtered traces recorded during sleep are shown in blue and during walking are shown in red; bandpassed signals are shown on top of each epoch in black, and the reference trace immediately below it, with detected oscillations in that frequency band indicated by a bold line. Traces from an alternate, simultaneously recorded site are plotted below the other trace. **c)**The temporal overlap of detected oscillations across brain sites, expressed as rate of concurrence. **d)**Likelihood of the behavior from the presence of an oscillation, shown on left plots; predictability is shown on the right, as the normalized likelihood of observing the behavior given the oscillatory bouts, factoring by the prior probability of the behavior. 90% bootstrapped confidence intervals are shown by errorbars. **e)**Polar histograms of preferred theta (5–10 Hz, black) and delta (1–5 Hz, gray) phase of multi-unit activity to local field potentials of hippocampal contacts for Animal 2 are shown. The MUA was tuned to local fields of both frequency bands but at different phases (p<0.001, Rayleigh test) suggesting that LFPs recorded in the hippocampus were not volume conducted.

Quantifying this inverse likelihood showed that no behavioral state is strongly or uniquely predicted by a given oscillatory band, however, oscillations at low frequencies best predict early-stage sleep, and oscillations at high frequencies best predict walking and other awake behaviors (Fig. 2d). This result is in striking contrast to findings in rodents, where theta oscillations are reliably observed in hippocampus during active behaviors such as walking rather than as an animal enters non-REM sleep ^1,19^.

In the primate, power spectral changes with behavioral state were generally similar across hippocampal and neocortical recordings sites, which could indicate volume conduction. If so, there should be temporal overlap of oscillatory bouts across sites, expressed as rate of concurrence across sites for a given frequency of oscillation. We found only around 20-50% concurrence between hippocampal contacts and other recording sites (Fig. 2c). That is, ~30% of the time hippocampal oscillations co-occurred with other recording sites. Moreover, hippocampal multiunit activity was partially phase-locked to the theta rhythm (5–10Hz) (p<0.001, Rayleigh test; Fig. 2e) suggesting that the LFPs recorded by hippocampal electrodes were effective in modulating local populations. Also, neocortical oscillations were generally stronger than those in the hippocampus, and spectral peaks were generally similar but not identical. Together, these results suggest that hippocampal activity was not simply volume conducted from neocortex.

Overall, we found that neocortical and hippocampal activity shifts from low-frequency (theta) activity in early-stage non-REM sleep to high frequencies (beta and gamma) during increasingly active behaviors including walking. In stark contrast to rodent studies, hippocampal theta oscillations were uncommon during walking and other active behaviors of monkeys. Theta oscillations are thought to be important for active waking processes including the formation of episodic memory and navigation not only for rodents ^19–23^; but also for humans ^13,17,24–26^. In this study, however, we show that theta oscillations in non-human primates (*i*) were not sustained during waking periods, (*ii*) did not track the durations of waking behaviors, and (*iii*) predicted sleep instead of active states. Beta and gamma frequencies, on the other hand, were better predictors for active and walking behaviors, consistent with a role for hippocampal beta/gamma in learning and navigation ^11–14,27^. Many of the spiking and high-frequency events commonly associated with theta oscillations such as phase coding^26^, phase precession ^20,28^, and gamma-frequency shifts ^29^ may nevertheless be conserved. These events could still underlie preserved mechanisms during exploration, without the necessity of coupling to a sustained, band-limited low frequency oscillation ^30^. Furthermore, cognitive states such as those responsible for abstract comparisons, flexible planning and strategizing may be transient and not time-locked to the behavioral classes we measured, including ambulatory movements. Our results suggest that however these elaborate cognitive states are supported by hippocampal and neocortical populations in primates, they are likely operating in the midst of a distinct set of spectral background conditions.

