## Supplemental Methods for "Hippocampal and neocortical oscillations are tuned to behavioral state in freely-behaving macaques"

### Online Methods

***Subjects and surgical implantations***

All procedures were approved by local ethics and animal care authorities. Three 10 kg adult female macaque monkeys (Macaca mulatta) underwent surgeries for electrode implantation conducted under sterile conditions and with the animals maintained under approximately 2% isoflurane anesthesia. Implantation of the electrodes was guided by pre-operative MR images, aligned to fiducial markers, using the Brainsight system (Rogue Research Inc., Montreal, Quebec, Canada). Animal 2 was implanted with an array of two electrode bundles, each containing 4 independently depth-adjustable platinum/tungsten multicore tetrodes (96 micron diameter; Thomas Recordings, Giessen, Germany). Post-operatively, tetrodes were lowered into the CA1/2 and CA3/DG regions of the right hippocampus (Fig. 1a, flipped to left hemisphere and marked as yellow). Recording locations were verified through MR/CT coregistration (Talakoub et al. 2016) and functional characterization of brain structures during lowering, including the appearance of sharp wave ripples (SWRs~~)~~ with complex spiking cells as shown in previous publications (SWRs: (Leonard and Hoffman 2017; Leonard et al. 2015) unit activity: (Leonard et al. 2015; Hussin et al. 2018); MR/CT: Talakoub et al. 2016). Animals 1 and 3 were implanted with 12 flexible probes each with 16 contacts targeted for anterior and intermediate-posterior hippocampus (designated aHPC, pHPC), medial temporal lobe rhinal cortex (aMTL), retrosplenial/posterior cingulate cortex (PCC), and anterior cingulate cortex/ventromedial prefrontal cortex (mPFC). Locations of implanted electrodes were verified for Animal 3 using postmortem high resolution gadolinium-enhanced MR images that resolved the electrode tracks. All animals’ recording sites were registered onto a reference brain (Seidlitz et al. 2018) shown as a 3-D cutout in Fig 1a, and as representative slices selected to depict each major location targeted in 2 coordinates and flattened on the 3^rd^ (mPFC, aMTL/aHPC, pHPC, as coronal slices and PCC as a horizontal slice). For monkeys 1 and 3, each probe connected to one of three, 64-channel electrode interface boards (EIBs), to which the wireless recording device was attached; Animal 2 was retrofit from the chronic tetrode drive connectors to the wireless device via cables and an adapter board.

***Electrophysiological recordings***

Local field potentials (LFPs) were referenced to the cranium and sampled at 30 KHz with 8KHz low-pass filter using 64/128 channel wireless systems (Neuralynx, Inc., Bozeman, Montana, USA). Animal 1 was recorded using a 128 channel wireless recording device. Two of the three EIBs (128 channels) were recorded at each session. EIB selection for each session was permuted resulting in similar overall recording time for each board. LFP signals were visually inspected before the start of each recording session. Neural signals were stored on an onboard microSD card without further transmission to the computer to reduce the power consumption and to extend the battery life of the recording device. Animal 2 and Animal 3 were recorded using a 64 channel wireless acquisition device connected to one EIB. LFP signals transmitted to a Digital Lynx SX (Neuralynx, Inc., Bozeman, Montana, USA) and stored on the acquisition computer.

***Video Recording and Synchronization with Electrophysiological Recordings***

Video frames were captured using Logitech HD Pro C920 webcam at 15 frames per second. Each frame was time-stamped with millisecond precision by a custom-made program, written in C. The program interfaced with the neural recording system via Netcom (Neuralynx, Inc., Bozeman, Montana, USA), a protocol to communicate with the acquisition system. Neural activity and video frames were synchronized using the events sent to the acquisition system by the video recording program.

Animal behavior was classified into four categories from the video recordings: sleep (periods of immobility when the eyes were closed for more than 3 minutes), inactive (being groomed by another animal, eating, and drinking), active (foraging, attentive visual orienting to anything outside the animal’s housing area, self-grooming, and grooming another animal), and walking (more than 3 steps). All sleep periods included in our analysis occurred after the room lights had been shut off (daily at 8 pm) and only the first 30 minutes of a night’s sleep was included in our analysis, to select for non-REM sleep epochs. The start and end of each behavior was marked on the video using a custom-made program by raters who were blind to the corresponding neural activity. The onset/offset times of the behaviors in the videos designated by the blind raters were then used to segment the continuously recorded stream of neural activity into epochs.

***Spectral Density of Local Field Potentials***

To identify peaks in the frequency spectrum above the noise level, the LFP at every recorded target location, for each behavioral epoch was segmented into 1-s epochs and transformed to the Fourier domain. Each epoch was corrected for 1/f by fitting a $\alpha/{f^{\gamma}}$ function to spectrum of the entire session at 0.25 Hz resolution. Finally, spectra were averaged for each behavior over all sessions.

***Oscillation Detection***

First, periods contaminated with EMG artifacts and other transients were excluded from our analyses. These periods were identified by monitoring the power in high frequency bands (>200 Hz). Muscle artifact is a broadband signal with considerable power at high frequency (>200 Hz) band. Thus, a simple method to identify the muscle and other common artifacts is to measure excessive high frequency activity across simultaneously-recorded electrode channels. We conservatively excluded 500 ms of activity preceding and following the artifacts to ensure the results are not contaminated with muscle and other artifactual activity.

To detect oscillations we used a Gaussian wavelet, providing a good trade-off between temporal and frequency resolutions. The 5-cycle wavelet atoms emphasized oscillations of at least 3 cycles. Average and standard deviation of wavelet values were calculated for each channel over the session. Oscillations were identified when wavelet values exceed one standard deviation from the mean ($\mu+\delta$) for over three cycles. Next, oscillatory time intervals for each frequency band were merged if adjacent oscillations were less than 250 ms apart. Finally, we measured occupancy rate of frequency bands for each behavior by normalizing the total duration of oscillation for a given behavior with total duration of the behavior.

***Specificity of Neural Activity to Behaviors***

In this section, we investigate whether the behavioral state can be identified based on cortical oscillations or alternatively, if these oscillations are not specific to any behavioral state. The predictability (information) of a behavior is related to the conditional probability of the behavior given the occurrence of the oscillation. Because conditional probabilities could be biased toward more frequent behaviors, conditional probabilities (C) were normalized by the prior probability of the behavioral state,

$$C\left( O,B \right)=\frac{p\left( B\vee O \right)}{p\left( B \right)}$$

where B represents behavior and O is the frequency band of the oscillation. This value is related to mutual information between behavioral state and occurrence of LFP oscillations that quantifies certainty of predicting the animal behavior by observing a bout of oscillation at a particular brain region.

Conditional probabilities are empirically calculated from the entire data. We estimated the confidence interval of the calculated conditional probabilities by bootstrapping the oscillation durations for each behavioral category; that is, conditional probabilities of sample distributions were calculated by resampling the oscillatory durations with replacement. After 1000 repetitions, 90% confidence intervals were estimated from the distributions.

***Concurrent oscillations across brain regions***

A concern with measures of oscillations observed across simultaneously recorded brain regions is the possibility that common activity or volume conduction underlies the signals in question. Common and volume conducted activity occur simultaneously or with a short lag. Thus, the timing of event epochs across sites, expressed as rate of concurrence across sites for a given frequency band, indicates whether or not observed oscillations are generated globally, as expected for common and volume conducted signal. Rate of concurrence across sites for a given frequency band were measured as the ratio between the time events were overlapping across two sites over the total time that events occur on either site.

***Preferred Delta-Theta Phase of Unit Activity***

The analysis of analog multi-unit activity (MUA) followed the method reported by Supèr and Roelfsema^6^. Data from Animal 2 with platinum tungsten tetrodes showing sharp-wave ripples in the posterior bundle was analyzed in this section. The broadband signal was segmented around the onset of the trial, bandpass filtered in between 750 and 5000 Hz and rectified. The rectified signal was low pass filtered at 1 KHz. The relationship between the phase of the delta (1–5 Hz) and theta (5–10 Hz) bands and the multi-unit amplitude was calculated by bandpass filtering the LFP signal at delta-theta frequencies and obtaining the instantaneous phase using the Hilbert transform. The MUA signal was rectified, and the histogram of MUA amplitude exceeding two standard deviations of its mean was calculated for each 0.05$\pi$ phase bin of the LFP. We computed a Rayleigh test for non-uniformity of circular data.

**Tables**

Table 1. Total duration of recordings and durations by behavioral group are listed per animal, in minutes.

| Animal | Total recording time | | | Sleep | Inactive | | | Active | | Walking |
| --- | --- | --- | --- | --- | --- | --- | --- | --- | --- | --- |
| Animal 1 | | | 10313 | 836 | | 1708 | 5126 | | 716 | |
| Animal 2 | | 1784 | | 1020 | 295 | | | 421 | | 47 |
| Animal 3 | | 506 | | 2 | 85 | | | 27 | | 390 |

**Video legend**

Examples of video recording and neural activity recorded wirelessly across various behavioral states (i.e. Sleeping, Walking, drinking (‘Inactive’), and being groomed (‘Inactive’) from three macaques. Each frame shows captured video (top), broadband field potentials (middle), and spectral decomposition of those signals (bottom). **Middle panel)** 2 seconds of neural activity recorded from aMTL, aHPC, pHPC, and PCC in amber, red, blue, and green color respectively – brain regions and colors assigned to them are also described at top left corner of the panel. The center of the graph (time t=0) corresponds to the video frame shown on the top panel. **Bottom panel)** spectral analyses of neural recordings marked with gray box in the middle panel (-500ms to 0sec). Color of the bars and brain regions matches the colors and regions of the middle panel. Fourier transform of the neural activity are shown in left and middle bar graphs. Bar graph at the right shows z-scored amplitude for each frequency band (1-4 Hz, 4-10Hz, 10-30 Hz, 30-90 Hz and 90-200 Hz).
